# Molecular and evolutionary strategies of meiotic cheating by selfish centromeres

**DOI:** 10.1101/405068

**Authors:** Takashi Akera, Emily Trimm, Michael A. Lampson

## Abstract

Asymmetric division in female meiosis creates selective pressure favoring selfish centromeres that bias their transmission to the egg. This centromere drive can explain the paradoxical rapid evolution of both centromere DNA and centromere-binding proteins despite conserved centromere function. Here, we define a molecular pathway linking expanded centromeres to histone phosphorylation and recrui™ent of microtubule destabilizing factors in an intraspecific hybrid, leading to detachment of selfish centromeres from spindle microtubules that would direct them to the polar body. We also introduce a second hybrid model, exploiting centromere divergence between species, and show that winning centromeres in one hybrid become losers in the other. Our results indicate that increasing destabilizing activity is a general strategy for drive, but centromeres have evolved distinct strategies to increase that activity. Furthermore, we show that drive depends on slowing meiotic progression, suggesting that a weakened meiotic spindle checkpoint evolved as a mechanism to suppress selfish centromeres.

## INTRODUCTION

Genomes are vulnerable to selfish genetic elements, which increase in frequency by forming additional copies of themselves (e.g., transposons) or distorting the transmission ratios during meiosis (i.e., meiotic drive), and are neutral or harmful to the host (McLaughlin and Malik, 2017). In female meiosis, selfish elements violate Mendel’s Law of Segregation by preferentially segregating to the egg, which increases their transmission to the progeny (Chmátal et al., 2017; Pardo-Manuel de Villena and Sapienza, 2001) (Figures S1A and S1B). Because centromeres direct chromosome segregation, they are the genetic elements with the best opportunity to cheat the segregation process. Meiotic drive of selfish centromeres, or centromere drive, can explain the “centromere paradox”: rapid evolution of both centromere DNA sequences and genes encoding centromere-binding proteins despite highly conserved centromere function in chromosome segregation (Henikoff et al., 2001). The centromere drive theory is based on the idea that natural selection favors centromere DNA sequences that act selfishly in female meiosis. Fitness costs associated with drive, for example due to deleterious alleles linked to driving centromeres, would also select for alleles of centromere-binding proteins that suppress drive. Thus, centromere DNA sequences and centromere proteins continually evolve in conflict with each other, analogous to a molecular arms race between viruses and the immune system. Supporting this theory, expanded centromeric satellite repeats in monkeyflowers preferentially transmit through female meiosis with associated fitness costs in pollen viability (Fishman and Saunders, 2008). Expanded centromeres also drive in mice, but a fitness cost has not been reported (Iwata-Otsubo et al., 2017; Wu et al., 2018). However, this theory raises several fundamental questions: how do centromeres cheat at a molecular level, linking from centromere expansion to selfish behavior, and how can centromere drive be suppressed?

To address these questions, we need a system to examine cell biological mechanisms of centromere drive. We previously established a *Mus musculus* hybrid between a standard laboratory strain with larger centromeres (either CF-1 or C57BL/6J) and a wild-derived strain from an isolated population with smaller centromeres (CHPO) (Figure 1A). In this intraspecific hybrid system (hereafter refer to as CHPO hybrid), larger centromeres have 6- to 10-fold more centromeric minor satellite repetitive sequence, more CENP-A nucleosomes which specify the site of kinetochore assembly, and more kinetochore proteins (e.g. CENP-C and HEC1) relative to smaller centromeres (Chmátal et al., 2014; Iwata-Otsubo et al., 2017). Larger centromeres in the CHPO hybrid drive by preferentially orienting towards the egg side of the spindle, and our previous results suggest that larger centromeres detach from the cortical side to re-orient towards the egg side (Figure S1B). These findings raise the question of why larger centromeres, which build larger kinetochores, are more susceptible to detachment. Moreover, it is unclear whether findings in one hybrid model system represent a general strategy for selfish centromeres to cheat. The large divergence in centromere DNA sequences between species (Kipling et al., 1995; Narayanswami et al., 1992; Wong et al., 1990) suggests that genetic conflict between centromere DNA and centromere-binding proteins has generated different evolutionary trajectories in different species and potentially different mechanisms of centromere drive.

**Figure 1.**
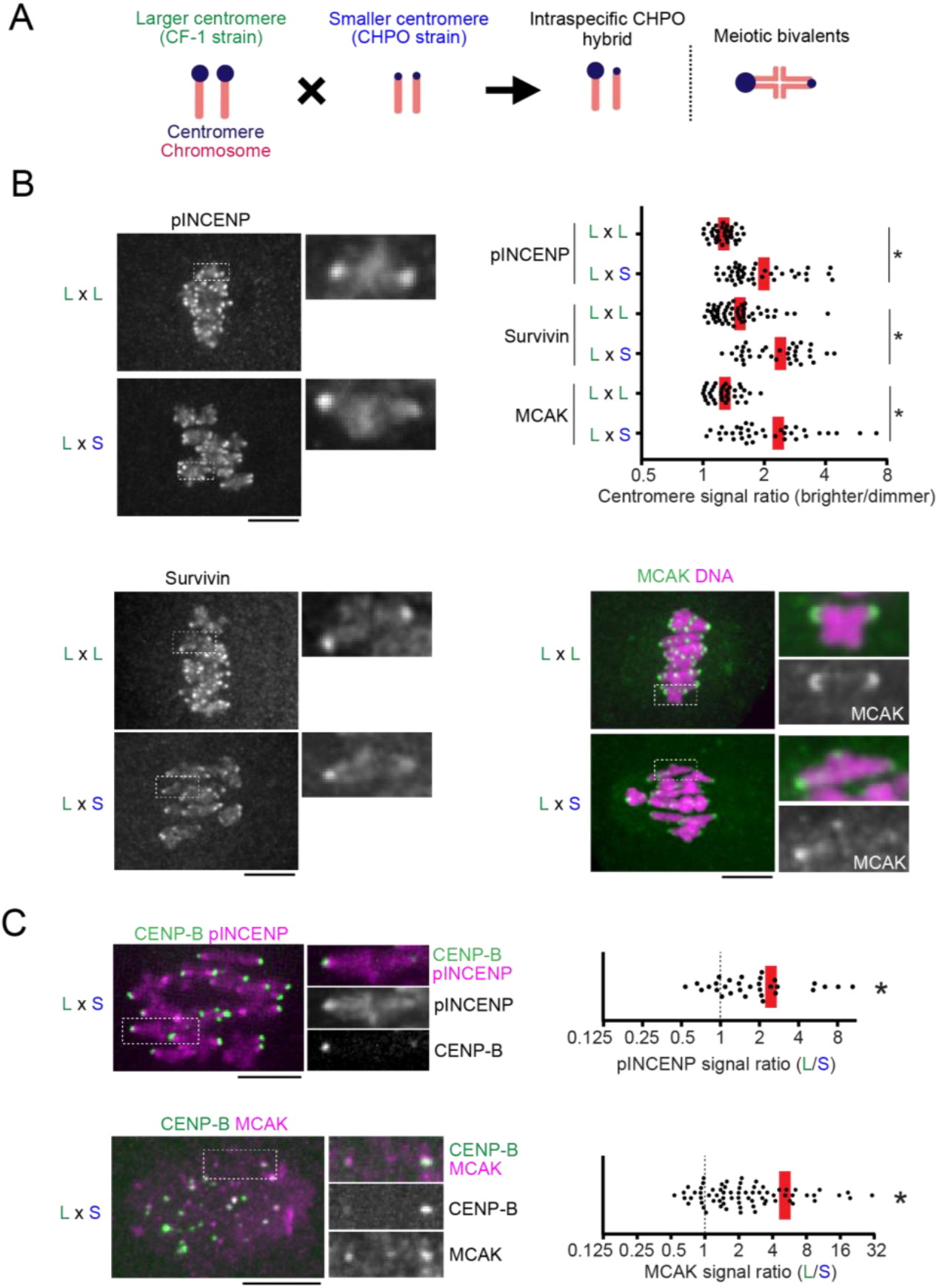
Selfish centromeres enrich more MT-destabilizing factors. (**A**) Schematic of the intraspecific CHPO hybrid system for centromere drive. A *Mus musculus* strain with larger (L) centromeres, CF-1, is crossed to a strain with smaller (S) centromeres, CHPO. In the hybrid offspring, chromosomes with larger and smaller centromeres are paired in meiotic bivalents. (**B**) CF-1 x CHPO (L x S) hybrid oocytes, or CF-1 x CF-1 (L x L) as controls, were fixed at metaphase I and stained for phosphorylated INCENP, Survivin, or MCAK. Graph shows centromere signal ratios, calculated as the brighter divided by the dimmer signal for each bivalent. (**C**) CF-1 x CHPO hybrid oocytes expressing CENP-B-EGFP were stained for pINCENP or MCAK. Graph shows centromere signal ratios, calculated as the CF-1 centromere divided by the CHPO centromere signal for each bivalent. Images (B, C) are maximum intensity z-projections showing all chromosomes (left), or optical slices magnified to show single bivalents (right); scale bars, 10 μm. In the graphs, each dot represents a single bivalent (n > 31 for each condition); red line, mean. \**P* < 0.001, indicating significant deviation from 1 in (C).

In this study, we uncovered molecular mechanisms and evolutionary strategies of meiotic cheating by selfish centromeres, exploiting both intraspecific variation and interspecific divergence in combination with cell biological analyses and experimental manipulation of centromeres. We establish a molecular pathway linking larger kinetochores to MT-destabilizing activity at peri-centromeres and show that this activity makes centromeres selfish. Moreover, we show that centromeres from different species have evolved distinct strategies to enrich destabilizing activity. Finally, our findings indicate that rapid progression through meiosis I is a mechanism to suppress drive.

## RESULTS

### BUB1 links larger kinetochores and higher MT-destabilizing activity at selfish centromeres

To confirm that bivalents in CHPO hybrid oocytes preferentially re-orient to direct larger centromeres towards the egg side during metaphase I, we live-imaged these flipping events. Since larger centromeres have expanded minor satellite repeats, we can distinguish larger and smaller centromeres by expressing fluorescently-tagged CENP-B protein, which binds minor satellite (Masumoto et al., 1989). We find that flipping events are biased to detach larger centromeres from the cortical side and re-orient them towards the egg side (Figure S1C). Consistent with this result, we previously showed that the orientation of larger centromeres is initially unbiased, but later becomes biased towards the egg side of the spindle just before anaphase I (Figure S1B). We also showed that larger centromeres form more unstable MT attachments compared to smaller centromeres, particularly with the cortical side of the spindle (Figure S1B, Early Meta I) (Akera et al., 2017). These findings suggest that selfish larger centromeres with larger kinetochores detach more readily from the spindle. To uncover the underlying mechanisms, we examined factors that destabilize MTs at centromeres to correct erroneous attachments: MCAK (mitotic centromere associated kinesin), which is a member of the kinesin-13 family, and the chromosome passenger complex (CPC) composed of Survivin, Borealin, INCENP, and Aurora B kinase (Carmena et al., 2012; Godek et al., 2015; Lampson and Grishchuk, 2017). By analyzing the bivalents in CHPO hybrid oocytes, we found asymmetry in MCAK, Survivin, and phosphorylated INCENP (Salimian et al., 2011) as a marker of active Aurora B kinase (Figure 1B). We did not observe such asymmetry in control oocytes in which centromeres of homologous chromosomes should be the same. Using CENP-B to label minor satellite, we found that larger centromeres have more of these MT-destabilizing factors compared to smaller centromeres (Figure 1C), similar to previous results for kinetochore proteins (Iwata-Otsubo et al., 2017). These observations suggest that selfish centromeres enrich MT-destabilizing activity to detach MTs and re-orient on the spindle.

MT-destabilizing factors localize to peri-centromeres, which are nearby but distinct chromosome regions from centromeres (Watanabe, 2012). Further, the amount of peri-centromeric repetitive major satellite DNA is similar between larger and smaller centromeres in the CHPO hybrid (Iwata-Otsubo et al., 2017). Therefore, it was unclear how larger centromeres can enrich more destabilizing activity. Shugoshin serves as a scaffold for both CPC and MCAK and is recruited to peri-centromeres by directly binding histone H2A threonine 121 phosphorylation marks (H2A pT121) (Watanabe, 2012). This histone phosphorylation is catalyzed by BUB1 kinase, which localizes at kinetochores (Kawashima et al., 2010). We hypothesized that larger centromeres recruit higher levels of BUB1 kinase relative to smaller centromeres, which would induce the asymmetric localization of Shugoshin and MT-destabilizing factors. Indeed, we found asymmetry in BUB1, H2A pT121, and SGO2, the major Shugoshin paralog in mouse oocytes (Lee et al., 2008), across the bivalents in CHPO hybrid oocytes (Figures 2A and S2) but not in control oocytes. Since MCAK is enriched more on larger centromeres relative to smaller centromeres (Figure 1C), co-staining with MCAK revealed that BUB1 and SGO2 are also higher on larger centromeres. Subsequently, co-staining of H2ApT121 with SGO2 showed that H2ApT121 is also higher on larger centromeres (Figure 2A). Together, these results indicate that BUB1 kinase is the molecular link between larger kinetochores and MT-destabilizing factors (Figure 2B).

**Figure 2.**
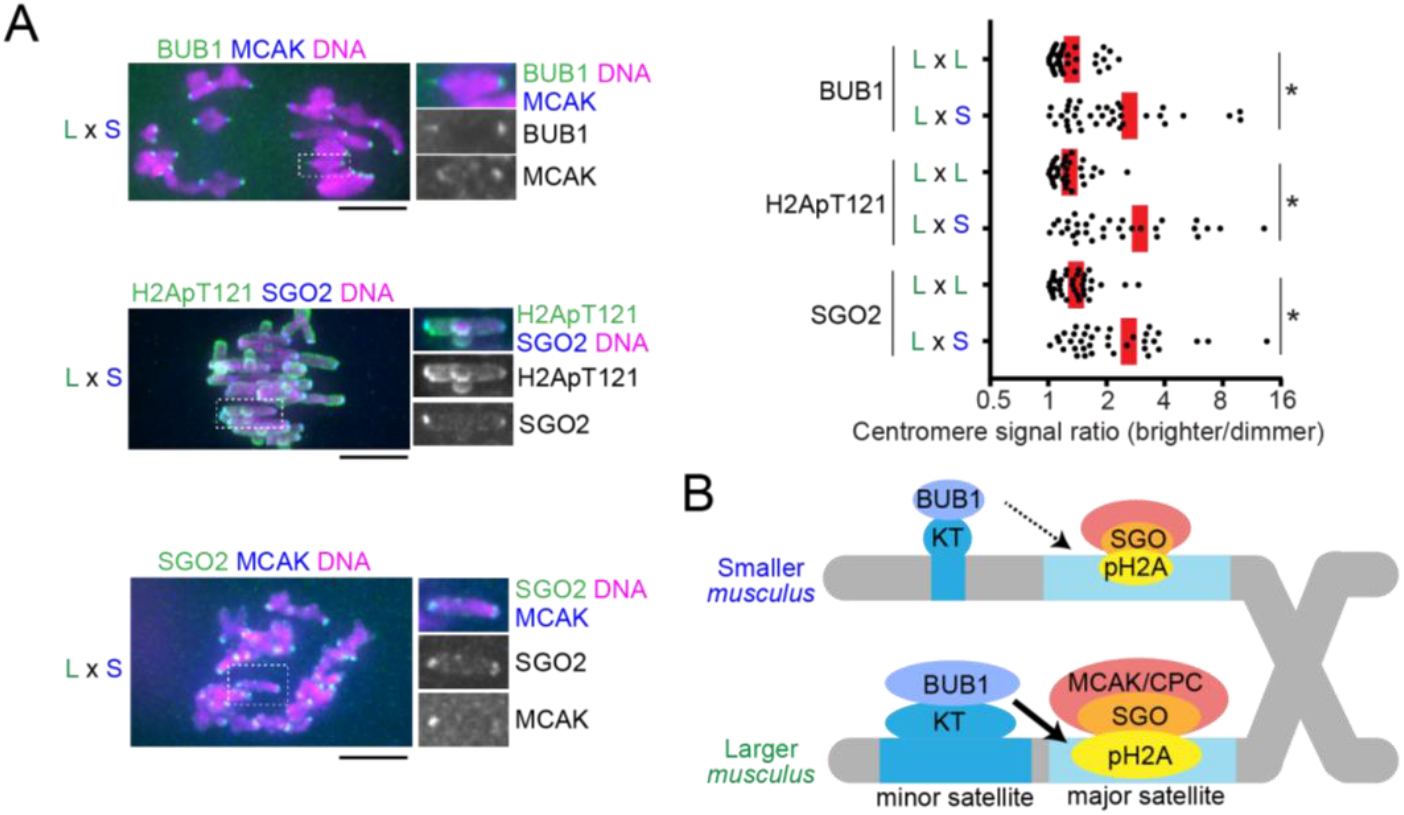
BUB1 links larger kinetochores and higher MT-destabilizing activity at selfish centromeres. (**A**) CF-1 x CHPO (L x S) hybrid oocytes, or CF-1 x CF-1 (L x L) as controls, were fixed at metaphase I and stained for BUB1, H2ApT121, or SGO2. Images are maximum intensity z-projections showing all chromosomes (left), or optical slices magnified to show single bivalents (right); scale bars, 10 μm. Graph shows centromere signal ratios, calculated as the brighter divided by the dimmer signal for each bivalent. Each dot represents a single bivalent (n > 32 for each condition); red line, mean; \**P* < 0.001. (**B**) Model of the amplified BUB1 pathway in larger centromeres compared to smaller centromeres in the intraspecific CHPO hybrid.

### Asymmetry in MT destabilizing activity is essential for centromere drive

To test the significance of the BUB1 pathway and MT-destabilizing activity for centromere drive, we developed an approach to experimentally equalize destabilizing activity between larger and smaller centromeres. We took advantage of the peri-centromeric repetitive major satellite DNA, which is similar between larger and smaller centromeres (Iwata-Otsubo et al., 2017), by genetically fusing the kinase domain of BUB1 to a TALE construct that targets major satellite (hereafter, Major Sat-BUB1) (Miyanari et al., 2013) (Figure 3A). Expressing this construct in hybrid oocytes increased MCAK and CPC levels on both sides of the bivalent and canceled the asymmetry (Figures 3B and S3). To determine the functional consequences of BUB1 targeting, we first examined the position of hybrid bivalents, which are off-center on the spindle at metaphase I in control hybrid oocytes, with larger centromeres closer to the pole (Figure S1B), indicating functional differences in microtubule (MT) interactions between larger and smaller centromeres (Chmátal et al., 2014). Bivalents were positioned close to the equator in hybrid oocytes expressing Major Sat-BUB1 (Figure 3C), which strongly suggests that centromere functions are equalized by this manipulation. Second, we confirmed that increasing MT-destabilizing factors at centromeres through BUB1 targeting indeed makes MTs more unstable based on cold-stable kinetochore-MT fibers (Rieder, 1981) (Figure 3D). Finally, we measured the orientation of hybrid bivalents on the spindle, using CENP-B to distinguish larger and smaller centromeres. We found that larger centromeres are no longer biased towards the egg pole in oocytes expressing Major Sat-BUB1, demonstrating that the asymmetry in destabilizing activity is essential for centromere drive (Figure 3E). Together, these results indicate that selfish centromeres in the intraspecific CHPO hybrid exploit BUB1 signaling at kinetochores to recruit MT destabilizers to win the competition in female meiosis.

**Figure 3.**
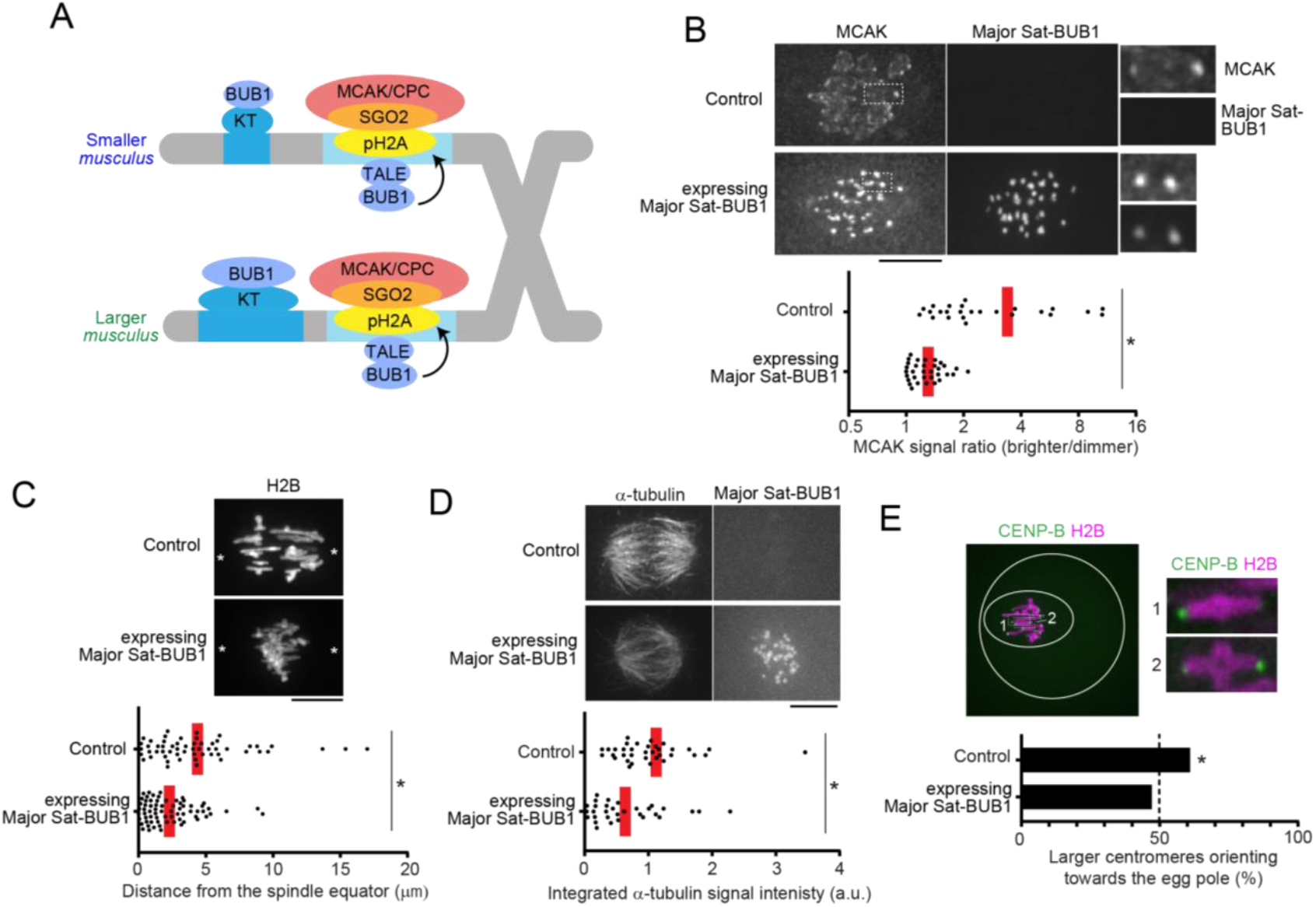
Asymmetry in MT destabilizing activity is essential for centromere drive. (**A**) Schematic of the strategy to equalize MT-destabilizing activity between larger and smaller centromeres by targeting BUB1 to major satellite. (**B**) CF-1 x CHPO (L x S) oocytes expressing a TALE targeting major satellite fused to the fluorescent protein mClover and to BUB1 lacking the N-terminal kinetochore-targeting domain (Major Sat-BUB1). Cells were fixed at metaphase I and stained for MCAK. Graph shows centromere signal ratios, calculated as the brighter divided by the dimmer signal for each bivalent. Each dot represents a single bivalent (n > 25 for each condition); red line, mean. (**C**) CF-1 x CHPO oocytes expressing Major Sat-BUB1 and H2B-EGFP were imaged live at metaphase I. Asterisks indicate the position of spindle poles determined by differential interference contrast imaging. Graph shows the distance between the spindle equator and the crossover position of each bivalent (n > 60 bivalents for each condition). (**D**) CF-1 x CHPO oocytes expressing Major Sat-BUB1 were analyzed for cold-stable MTs at metaphase I. Graph shows integrated α-tubulin signal intensity in the spindle (n > 32 spindles for each condition). (**E**) CF-1 x CHPO oocytes expressing Major Sat-BUB1, CENP-B-mCherry, and H2B-EGFP were imaged live shortly before anaphase I onset (within 30 min). The fraction of bivalents with the larger centromere oriented towards the egg pole was quantified (n > 106 bivalents for each condition). Images (B-E) are maximum intensity z-projections or optical slices magnified to show single bivalents. \**P* < 0.005, indicating significant deviation from 50% in (E).

### Centromeres in an interspecific hybrid exhibit asymmetry in destabilizers but not in kinetochore size

Our experiments with the intraspecific CHPO hybrid system revealed molecular mechanisms of drive, linking selfish centromeres to increased kinetochore size and recrui™ent of MT-destabilizing factors to the peri-centromere. To determine whether these findings represent general properties of driving centromeres, we exploited the large divergence in centromere DNA sequences between mouse species (Wong et al., 1990) to identify a second hybrid model for centromere drive. We selected *Mus musculus* and *Mus spretus* because centromere DNA has diverged between the two species with *spretus* centromeres having substantially more minor satellite and less major satellite repeats compared to *musculus* centromeres (Hale et al., 1993; Wong et al., 1990). Therefore, we crossed SPRET/EiJ (*Mus spretus*) with CF-1 or C57BL/6J (*Mus musculus* with larger centromeres relative to CHPO) to produce an interspecific hybrid (hereafter, *spretus* hybrid) (Figure 4A). We used CF-1 to mate with CHPO in the intraspecific cross because they efficiently produce hybrid offspring, but we primarily used C57BL/6J as the *musculus* strain in the *spretus* cross because of difficulties mating *spretus* with CF-1.

**Figure 4.**
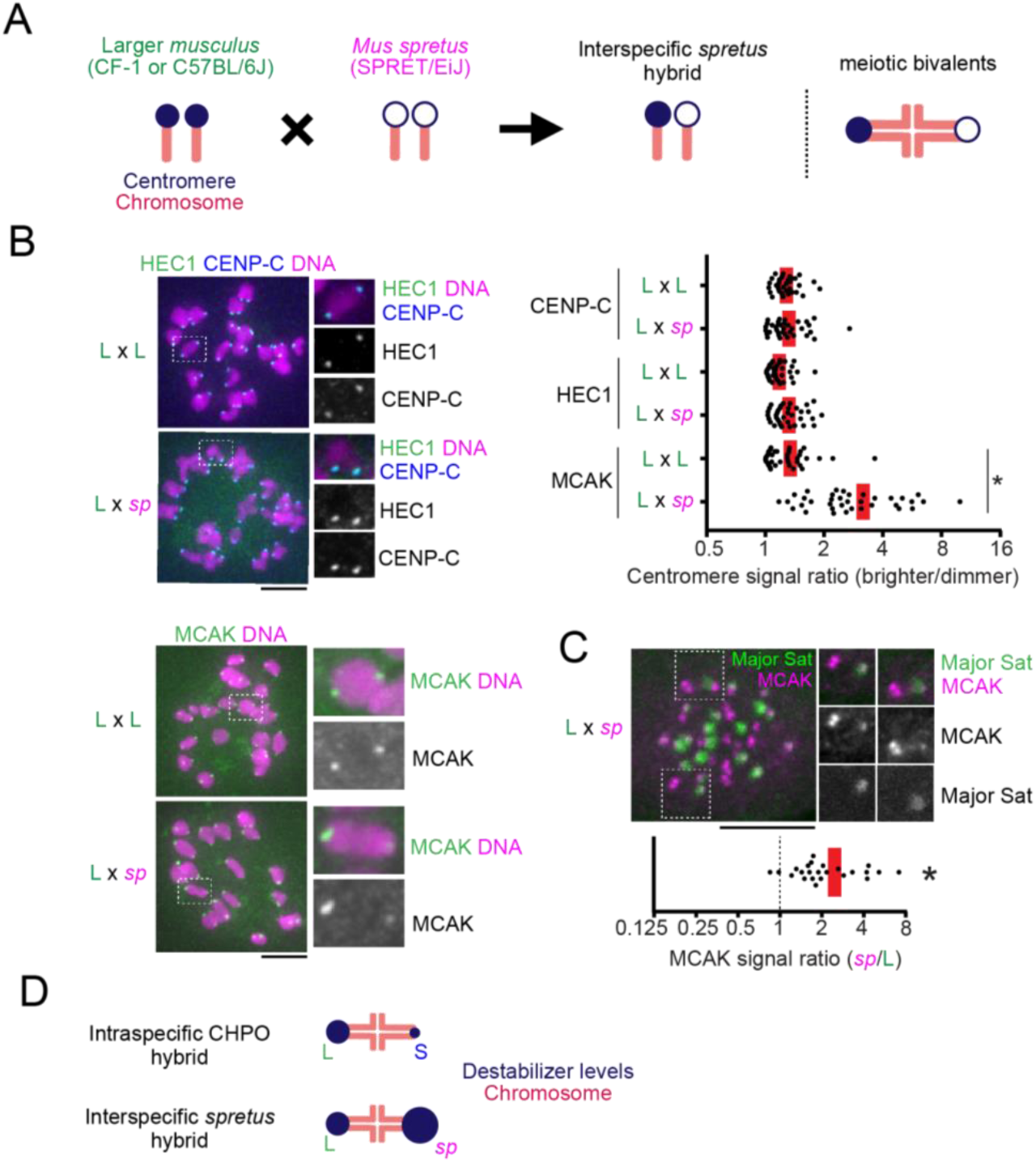
Centromeres in an interspecific hybrid exhibit asymmetry in destabilizers but not in kinetochore size. (**A**) Schematic of the interspecific *spretus* hybrid system. A *Mus musculus* strain with larger centromeres (L, CF-1 or C57BL/6J) is crossed to a *Mus spretus* strain (*sp*, SPRET/EiJ). In the hybrid offspring, chromosomes with *musculus* and *spretus* centromeres are paired in meiotic bivalents. (**B**) C57BL/6J x SPRET/EiJ (L x *sp*) hybrid oocytes, or C57BL/6J x C57BL/6J (L x L) as controls, were fixed at metaphase I and stained for the indicated centromere proteins. Graph shows centromere signal ratios, calculated as the brighter divided by the dimmer signal for each bivalent (n > 36 bivalents for each condition). (**C**) C57BL/6J x SPRET/EiJ (L x *sp*) oocytes expressing Major Sat. TALE-mClover were stained for MCAK. Graph shows centromere signal ratios, calculated as the *spretus* centromere divided by the C57BL/6J centromere signal for each bivalent (n = 24 bivalents). Images (B, C) are maximum intensity z-projections showing all chromosomes (left), or optical slices magnified to show single bivalents (right); scale bars, 10 μm. In the graphs, each dot represents a single bivalent; red line, mean. \**P* < 0.001, indicating significant deviation from 1 in (C). (**D**) Schematic of relative MT destabilizer levels in both intraspecific and interspecific hybrid models.

We first measured centromere protein levels in *spretus* hybrid oocytes. Both the inner kinetochore protein CENP-C that binds to CENP-A nucleosomes and the outer kinetochore protein HEC1 that binds to MTs were similar across the hybrid bivalents (Figure 4B). In contrast, MCAK showed significant asymmetry across the bivalents in the *spretus* hybrid, but not in control *musculus* oocytes (Figure 4B). The CPC localized all over the chromosomes in this hybrid without obvious enrichment on centromeres (Figure S4), probably due to higher levels of cohesin complex on chromosome arms (Sodek et al., 2017), which contributes to CPC recrui™ent through the Haspin kinase pathway (Goto et al., 2017). Therefore, we focused on MCAK as a MT-destabilizing factor in the *spretus* hybrid. Since *musculus* centromeres have more peri-centromeric repetitive major satellite DNA, we can distinguish *musculus* and *spretus* centromeres by expressing a fluorescently-tagged TALE construct that recognizes major satellite (Miyanari et al., 2013). Using this approach, we found that the larger *musculus* centromeres, which recruited more MCAK in the intraspecific CHPO hybrid, have less MCAK compared to *spretus* centromeres in the interspecific hybrid (Figures 1C, 4C, and 4D).

We performed two assays to test the functional consequences of MCAK asymmetry. First, we predicted that centromeres with more MCAK should initiate flipping events by detaching MTs first. To test this prediction, we tracked the flipping process by live-imaging to determine which centromere moves towards the opposite pole first to initiate flipping, indicating that it detaches first. We found that larger *musculus* centromeres initiated 76% of flipping events in the intraspecific CHPO hybrid (Figure 5A), whereas the same *musculus* centromeres initiated only 25% of flipping in the interspecific *spretus* hybrid (Figure 5B). These results are consistent with relative MCAK levels: *spretus* > larger *musculus* > smaller *musculus* (Figures 1C and 4C). Second, based on the findings from the CHPO hybrid system, we predicted that *spretus* hybrid bivalents would be positioned off-center on the spindle, with centromeres with higher destabilizing activity closer to the pole. Indeed, we found that *spretus* centromeres with more MCAK are closer to the pole (Figure 6A). Together, these results establish an interspecific hybrid system in which *spretus* centromeres have higher destabilizing activity independent of kinetochore size.

**Figure 5.**
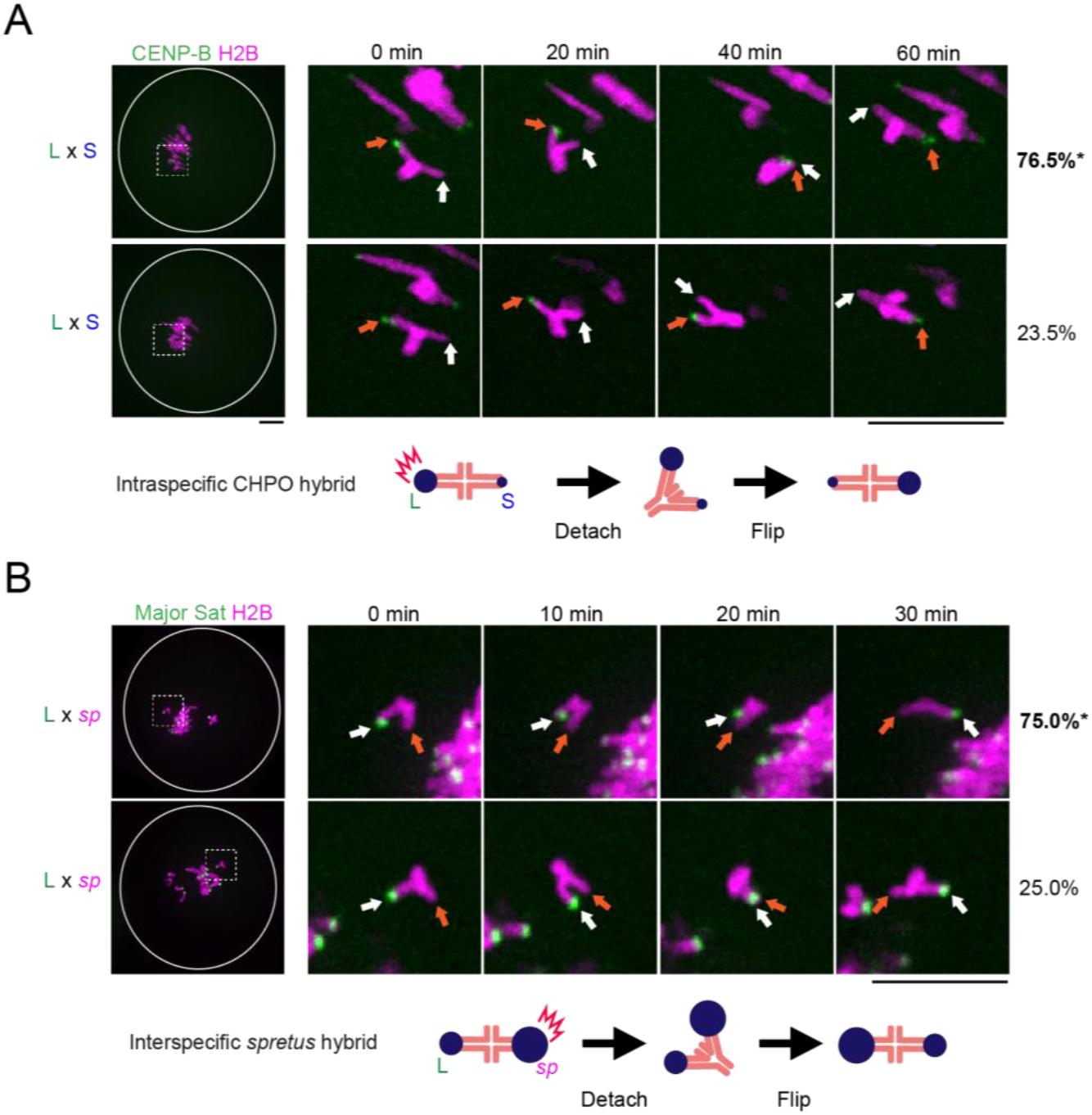
Relative MCAK levels on centromeres predict their destabilizing activity in flipping events. (**A**) CF-1 x CHPO (L x S) oocytes expressing CENP-B-mCherry and H2B-EGFP were imaged live. Time-lapse images show examples of flipping events, which were analyzed to determine the frequency of either the larger (orange arrows) or smaller (white arrows) *musculus* centromere moving first to initiate flipping (top and bottom panels respectively). Percentages on the right indicate frequency of each case (n = 45 flipping events from 61 cells). (**B**) CF-1 x SPRET/EiJ and C57BL/6J x SPRET/EiJ (L x *sp*) oocytes expressing Major Sat. TALE-mClover and H2B-mCherry were imaged live and analyzed to determine whether the *spretus* (orange arrows) or larger *musculus* (white arrows) centromere initiates flipping (top and bottom panels respectively) (n = 27 flipping events from 20 cells). Images (A, B) are maximum intensity z-projections showing all chromosomes (left), or optical slices magnified to show flipping events (timelapse). White circle indicates the cell outline. Schematics show the more frequent flipping events, with relative MCAK levels indicated by the size of the blue circle. Scale bars, 10 μm. \**P* < 0.05, indicating significant deviation from 50%.

**Figure 6.**
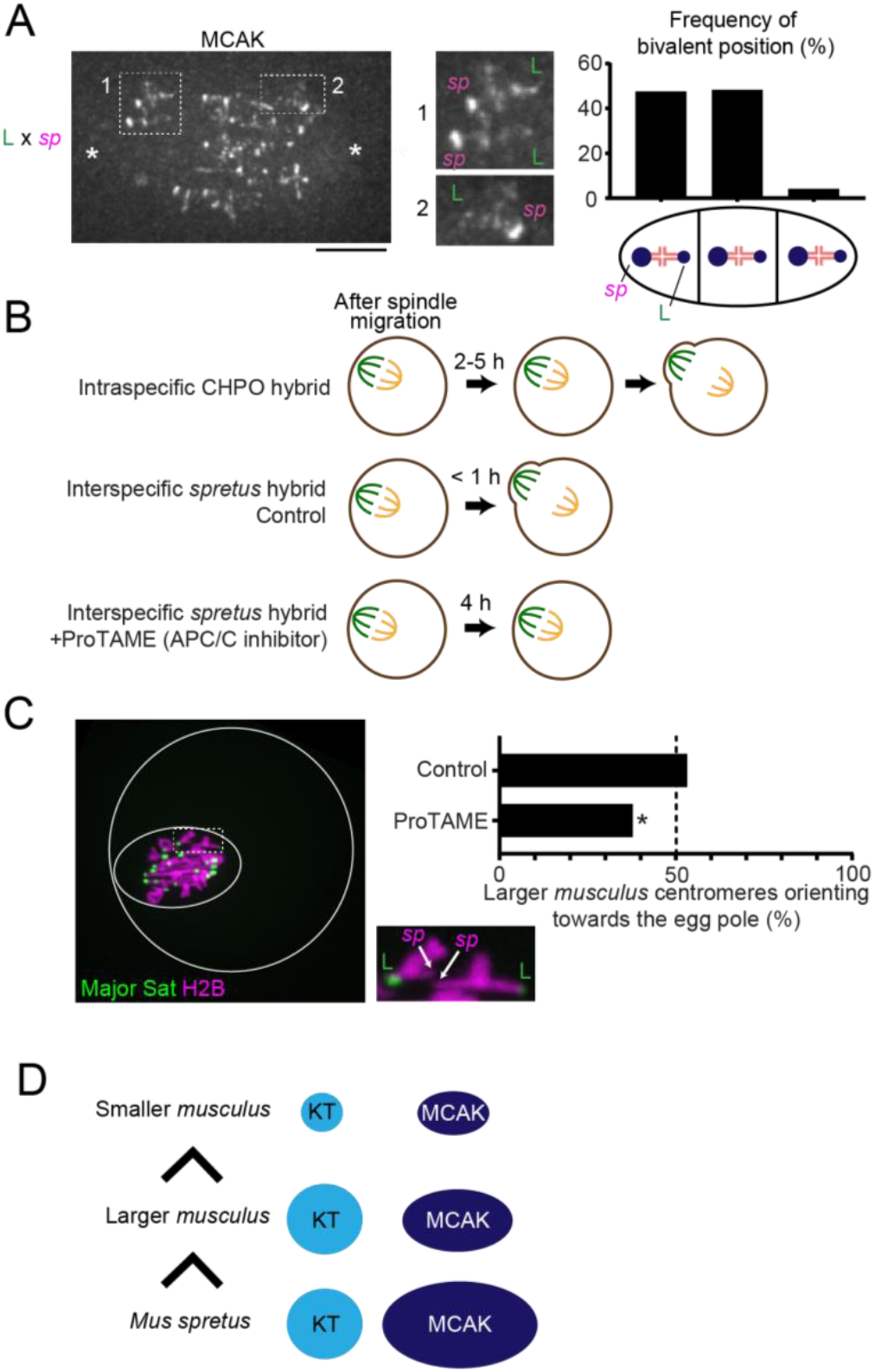
Relative MT-destabilizing activity determines the direction of centromere drive. (**A**) C57BL/6J x SPRET/EiJ (L x *sp*) oocytes were fixed at metaphase I and stained for MCAK. Images are maximum intensity z-projections showing all chromosomes or optical slices magnified to show two bivalents closer to the left pole (1) or a single bivalent closer to the right pole (2). Schematic shows bivalent positions as equidistant between the two poles (middle) or off-center with either the *spretus* centromere or the larger *musculus* centromere closer to the pole. The frequency of each case is plotted (n = 120 bivalents). (**B**) Schematics of meiotic progression. ProTAME trea™ent allows the *spretus* hybrid to delay anaphase I onset at least 4 hours, comparable to the CHPO hybrid. (**C**) CF-1 x SPRET/EiJ and C57BL/6J x SPRET/EiJ oocytes expressing Major Sat. TALE-mClover and H2B-mCherry were imaged live either shortly before anaphase I (control) or 4 hours after spindle migration (ProTAME). Images are a maximum intensity z-projection of the whole oocyte (left) and an optical slice magnified to show two bivalents (right). Solid and dashed white circles indicate the outline of the cell and the spindle, respectively. Graph shows the fraction of bivalents with the larger *musculus* centromere oriented towards the egg pole (n > 135 bivalents for each condition). \**P* < 0.005, indicating significant deviation from 50%. Scale bars, 10 μm. (**D**) Schematic showing that the direction of centromere drive correlates with MT-destabilizer levels but not with kinetochore size.

### Winning centromeres in one hybrid become losers in another hybrid based on the relative destabilizing activity

If higher destabilizing activity is a general property of driving centromeres, the prediction is that larger *musculus* centromeres, which win in the CHPO hybrid by preferentially orienting towards the egg side of the spindle, would be losers in the *spretus* hybrid. We found that the orientation of *spretus* hybrid bivalents was unbiased just before anaphase I (Figure 6C, Control), but we also noticed that *spretus* hybrid oocytes do not delay anaphase onset, in sharp contrast to the CHPO hybrid (Akera et al., 2017; Sebestova et al., 2012) (Figure 6B). Timing is important because the spindle initially forms in the center of the oocyte, and later migrates towards the cortex (Almonacid et al., 2014; Azoury et al., 2011; Holubcová et al., 2013). CDC42 signaling from the cortex regulates MT tyrosination, which generates asymmetry between the two sides of the spindle after spindle migration (Akera et al., 2017; Dehapiot et al., 2013) (Figures 6B, S1, and S5). Biased orientation arises from biased flipping while the spindle is positioned close to the cortex and asymmetric (Figures S1B and S1C). Consistent with this idea, the orientation of CHPO hybrid bivalents is initially unbiased at the earlier stage right after spindle migration, but anaphase I is delayed 2 - 5 hours, which provides time for the flipping events (Akera et al., 2017) (Figure S1B). In contrast, *spretus* hybrid oocytes progress to anaphase I immediately after spindle migration.

These results suggest that centromere drive depends on slowing meiotic progression so that selfish centromeres have time to flip after the spindle has acquired asymmetry. Therefore, we experimentally delayed anaphase in the *spretus* hybrid using an Anaphase Promoting Complex/Cyclosome (APC/C) inhibitor, ProTAME (Zeng et al., 2010) (Figure 6B). This delay induced biased orientation 4 hours after spindle migration, with larger *musculus* centromeres preferentially oriented towards the cortical side of the spindle, which will direct them to the polar body (Figure 6C). Together, these results demonstrate that relative destabilizing activity defines the directionality of centromere drive and that centromere drive depends on slowing meiotic progression.

## DISCUSSION

Our findings reveal both molecular and evolutionary strategies of meiotic cheating by selfish centromeres. Our results from the intraspecific CHPO hybrid model are consistent with the centromere drive theory, which proposes that selfish centromeres expand centromeric satellite repeats and build larger kinetochores to win the competition in female meiosis (Henikoff et al., 2001; Iwata-Otsubo et al., 2017). However, it has been unclear how larger kinetochores lead to selfish behavior. Centromeres incorporate both MT-binding activity at kinetochores and counteracting MT-destabilizing activities, which promote re-orientation of incorrect attachments to prevent segregation errors (Heald and Khodjakov, 2015). We show that selfish centromeres exploit the same destabilizing activity to bias their segregation to the egg. Multiple lines of evidence support this conclusion. First, we observed higher levels of MT-destabilizing factors at selfish centromeres in both intraspecific and interspecific hybrids (Figures 1 and 4). Second, equalizing destabilizers across hybrid bivalents prevented drive (Figure 3). Third, selfish centromeres initiated flipping events by detaching MTs (Figure 5). Fourth, relative levels of MT-destabilizing factors determine the direction of centromere drive, converting winners in one hybrid to losers in another hybrid (Figure 6).

Our finding that MT-destabilizing activity underlies non-Mendelian segregation is consistent with the cell biology of chromosome segregation in mouse oocytes (Kitajima, 2018). Initial MT attachments are established when the spindle is still symmetric and therefore lacks spatial cues to guide selfish centromeres (Kitajima et al., 2011), which must selectively detach to flip towards the egg pole after the spindle has migrated and acquired asymmetry. This process implies some destabilizing activity that acts specifically on the cortical side of the spindle, which is more tyrosinated (Akera et al., 2017). We propose MCAK as this activity because it preferentially destabilizes tyrosinated MTs (Peris et al., 2009; Sirajuddin et al., 2014) and is recruited at higher levels to selfish centromeres in both hybrid models. Also, MCAK localizes to centromeres only at late metaphase I (Illingworth et al., 2010), which matches the timing of flipping to orient selfish centromeres towards the egg pole.

We show that increased kinetochore size is one mechanism linking expanded centromeres to recrui™ent of MT-destabilizing factors at the peri-centromere, and we establish the underlying molecular pathway. In the intraspecific CHPO hybrid, larger kinetochores lead to more BUB1 kinase and histone phosphorylation, which recruits Shugoshin and MT-destabilizing factors (Figure 2). By experimentally equalizing destabilizing activity through BUB1 targeting to major satellite sequences, we demonstrate the significance of this pathway for centromere drive in this *Mus musculus* hybrid (Figure 3). However, *Mus spretus* centromeres do not depend on increased kinetochore size to enrich destabilizing activity. These results suggest that genetic conflict between centromere DNA and centromere-binding proteins has played out differently in different species, leading to distinct mechanisms to enrich the same activity to bias segregation. At the level of centromere DNA, we find that more expanded centromeres win in female meiosis, consistent with the drive theory (Henikoff et al., 2001): larger *musculus* centromeres win against smaller *musculus* centromeres but lose against *spretus* centromeres, which have even more minor satellite DNA (Figure 6D). How *spretus* centromeres enrich high destabilizing activity without building bigger kinetochores is an important future question, especially because *spretus* centromeres have very little major satellite, which is the major site in *Mus musculus* for peri-centromeric heterochromatin (Guenatri et al., 2004).

The core of the centromere drive theory is the idea that suppression of drive provides selective pressure for evolution of centromere proteins (Henikoff et al., 2001). Although this idea has been influential to explain the paradoxical rapid evolution of centromere proteins, it has been difficult to directly test without some understanding of the cell biological basis of centromere drive. Our results provide the first step towards a mechanistic model for the selective pressure. We show that flipping events to face selfish centromeres towards the egg pole take time, and rapid progression through meiosis I prevents drive (Figures 6B and 6C). Meiotic progression is controlled by the spindle assembly checkpoint, which delays anaphase until all chromosomes are attached to spindle MTs (Joglekar, 2016). This checkpoint is weaker in oocytes compared to somatic cells, which is counter-intuitive because of the risk of producing aneuploid eggs (Nagaoka et al., 2011; Shao et al., 2013). We propose that the weakened checkpoint may be adaptive by suppressing drive of selfish centromeres. Multiple mechanisms could weaken the spindle assembly checkpoint, for example dampening the signaling cascade at centromeres or strengthening APC/C activity. Moreover, a large cytoplasmic volume, which is a general feature of female meiosis, is directly linked to the weakened checkpoint (Kyogoku and Kitajima, 2017; Lane and Jones, 2017). Identifying genes with signature of rapid evolution may provide further insights into how genomes have evolved to suppress drive, through either a weak checkpoint or other mechanisms. Our cell biological studies of centromere drive, combined with such molecular evolution analysis, will lead to a deeper understanding of the molecular arms race between selfish elements and the rest of the genome.

## Acknowledgements

We thank B. E. Black, R. M. Schultz, and M. T. Levine for comments on the manuscript, D. A. Compton for the MCAK antibody, Y. Watanabe for the CENP-C, SGO2, and BUB1 antibodies, and M. E. Torres-Padilla and Y. Miyanari for the TALE constructs. The research was supported by NIH grant GM122475 (M.A.L.) and by a research fellowship from Uehara Memorial Foundation (T.A.).

## STAR METHODS

### CONTACT FOR REAGENT AND RESOURCE SHARING

Further information and requests for resources and reagents should be directed to and will be fulfilled by the Lead Contact, Michael A. Lampson.

### EXPERIMENTAL MODEL AND SUBJECT DETAILS

#### Mice

Mouse strains were purchased from the Jackson Laboratory (ZALENDE/EiJ, stock #001392 corresponds to CHPO; C57BL/6J, stock# 000664; SPRET/EiJ, stock# 001146) and from Envigo (NSA, stock# 033 corresponds to CF-1). All mice used in this study were 8-14 week-old females. All animal experiments were approved by the Institutional Animal Care and Use Committee and were consistent with the National Institutes of Health guidelines.

### METHOD DETAILS

#### Oocyte collection and culture

Female mice were hormonally primed with 5U of Pregnant Mare Serum Gonadotropin (PMSG, Calbiochem, cat# 367222) 44-48 h prior to oocyte collection. Germinal vesicle (GV)-intact oocytes were collected in bicarbonate-free minimal essential medium with polyvinylpyrrolidone and Hepes (MEM-PVP) (Stein and Schindler, 2011), denuded from cumulus cells, and cultured in Chatot-Ziomek-Bavister (CZB) (Chatot et al., 1989) medium covered with mineral oil (Sigma, cat# M5310) in a humidified a™osphere of 5% CO2 in air at 37°C. During collection, meiotic resumption was inhibited by addition of 2.5 μM milrinone. Milrinone was subsequently washed out to allow meiotic resumption. Oocytes were checked for GVBD 1.5 h after milrinone washout, and those that did not enter GVBD stage were removed from the culture.

#### Oocyte microinjection

GV oocytes were microinjected with ∼5 pl of cRNAs in MEM-PVP containing milrinone at room temperature (RT) with a micromanipulator TransferMan NK 2 (Eppendorf) and picoinjector (Medical Systems Corp.). After the injection, oocytes were kept in milrinone for 16 h to allow protein expression. cRNAs used for microinjections were *H2B-mCherry* (human histone H2B with mCherry at the C-terminus) at 400 ng/μl, *TALE-mClover* (TALE construct that recognize Major satellite repeats fused to mClover and 3 tandem Halo tag at the C-terminus) at 1000 ng/μl, *H2B-Egfp* (human histone H2B with EGFP at the C-terminus) at 600 ng/μl, *Cenpb-mCherry* (mouse CENP-B with mCherry at the C-terminus) at 1300 ng/μl, and *Major Sat-Bub1* (TALE construct that recognize Major satellite repeats fused to mClover and the kinase domain of mouse BUB1 (a.a. 672-1058) at the C-terminus) at 100 ng/μl. cRNAs were synthesized using the T7 mScript™ Standard mRNA Production System (CELL SCRIPT).

#### Live imaging

Oocytes were placed into 2 μl drops of CZB media covered with mineral oil in a glass-bottom tissue culture dish (FluoroDish FD35-100) in a heated environmental chamber with a stage top incubator (Incubator BL and Heating Insert P; PeCon GmBH) to maintain 5% CO2 in air and 37°C. Confocal images were collected with a microscope (DMI4000 B; Leica) equipped with a 63× 1.3 NA glycerol-immersion objective lens, an xy piezo Z stage (Applied Scientific Instrumentation), a spinning disk confocal scanner (Yokogawa Corporation of America), an electron multiplier charge-coupled device camera (ImageEM C9100-13; Hamamatsu Photonics), and an LMM5 laser merge module with 488- and 593-nm diode lasers (Spectral Applied Research) controlled by MetaMorph software (Molecular Devices). Confocal images were collected as z-stacks at 1 μm intervals to visualize the entire meiotic spindle.

#### Oocyte immunocytochemistry

MI oocytes at different times after GVBD were cultured in CZB media. Oocytes were fixed in freshly prepared 2% paraformaldehyde in PBS with 0.1% Triton X-100, pH 7.4, for 20 min at RT, permeabilized in PBS with 0.1% Triton X-100 for 15 min at RT, placed in blocking solution (PBS containing 0.3% BSA and 0.01% Tween-20) overnight at 4 °C, incubated 1 h with primary antibodies in blocking solution, washed 3 times for 15 min, incubated 1 h with secondary antibodies, washed 3 times for 15 min, and mounted in Vectashield (Vector, cat# H-1400) with bisbenzimide (Hoechst 33342, Sigma-Aldrich) to visualize chromosomes. For Figure S5, 0.05% glutaraldehyde was added to the fixative to better preserve spindle MTs (Schuh and Ellenberg, 2007). The primary antibodies used for this study were rat anti-tyrosinated α-tubulin (1:1000, Serotec, YL1/2), rabbit anti-β-tubulin (9F3) monoclonal conjugated to Alexa Fluor 488 (1:50 dilution; Cell Signaling Technology), mouse anti-α-tubulin (1:500, Sigma, DM1A), CREST human autoantibody against centromere (1:100, Immunovision), rabbit anti-human p-INCENP (Salimian et al., 2011) (1:200), rabbit anti-human Survivin (1:500, Cell signaling, 71G4B7), rabbit anti-human MCAK (1:1000, a gift from Duane Compton), mouse anti-mouse BUB1 (1:100, a gift from Yoshinori Watanabe), mouse anti-mouse SGO2 (1:500, a gift from Yoshinori Watanabe), rabbit anti-histone H2AT120ph (1:2500, Active motif, 39391), mouse anti-mouse HEC1 (1:100, Santa Cruz, C-11), rabbit anti-mouse CENP-C (1:2000, a gift from Yoshinori Watanabe). Secondary antibodies were Alexa Fluor 488–conjugated donkey anti-mouse or donkey anti-rabbit or Alexa Fluor 594–conjugated donkey anti-rat, donkey anti-rabbit, donkey anti-mouse or goat anti-human (1:500, Invitrogen). Confocal images were collected as z-stacks at 1 μm intervals to visualize the entire meiotic spindle, using the spinning disc confocal microscope described above.

To quantify centromere signal ratios, optical slices containing centromeres from the same bivalent were added to produce a sum projection using Fiji/ImageJ. Ellipses were drawn around the centromeres, and signal intensity was integrated over each ellipse after subtracting background, obtained by measuring the average intensity of a region near the centromeres. Ratios were obtained for each bivalent by dividing the intensity of the brighter centromere by that of the dimmer centromere unless otherwise specified in the figure legend.

#### Biased orientation assay

GV oocytes from CF-1 x CHPO (CHPO hybrid) or SPRET/EiJ x CF-1 and SPRET/EiJ x C57BL/6J (*spretus* hybrid) were collected and microinjected with cRNAs encoding CENP-B-mCh and H2B-EGFP (CHPO hybrid) or Major Satellite-mClover and H2B-mCh (*spretus* hybrid). Oocytes were induced for meiotic resumption by washing out milrinone. Live imaging was performed as described above, starting 10 h (CHPO hybrid) or 7 h (*spretus* hybrid, Control) after GVBD to capture the time just before anaphase onset. Images were taken every 30 min. *Spretus* hybrid oocytes arrested in metaphase I by ProTAME were imaged at 10 h after GVBD. The position of the spindle near the cortex was confirmed by differential interference contrast images, and the fraction of bivalents with the larger *musculus* centromere (CF-1 or C57BL/6J) oriented towards the egg was quantified, using CENP-B to distinguish CF-1 centromeres from CHPO or Major Satellite to distinguish *spretus* centromeres from CF-1 or C57BL/6J. The CHPO strain also contains seven Robertsonian fusions, each of which pairs with two CF-1 chromosomes in MI to form a trivalent (Chmátal et al., 2014), but we included only bivalents in our analyses to avoid complications of trivalents.

#### Flipping assay

Oocytes from CF-1 x CHPO (CHPO hybrid) or SPRET/EiJ x CF-1 and SPRET/EiJ x C57BL/6J (*spretus* hybrid) were imaged live as in the biased orientation assay except for starting 7 h (CHPO hybrid) or 4.5 h (*spretus* hybrid) after GVBD and taking images every 10 or 20 min. To measure the frequency of each centromere initiating the flipping, we only analyzed flipping events in which we captured the intermediate state (only one of the two centromeres moving towards the opposite pole). To measure biased flipping, oocytes with the spindle completely migrated towards the cortex (distance between the cortex and the center of the spindle < 25 μm) were analyzed.

#### Cold-stable MT assay

Oocytes were placed into ice cold MEM-PVP for 6 min before fixation and stained for α-tubulin. Confocal images were collected to visualize the entire meiotic spindle, using the spinning disc confocal microscope described above. To calculate tubulin signal intensity, ellipses were drawn around the spindle, and α-tubulin intensity was integrated over each ellipse in optical slices containing the spindle, after subtracting background.

#### Statistical analysis

Data points are pooled from at least two independent experiments unless specified in the figure legend. The following statistical methods were used: unpaired t test in Figures 1B, 2A, 3B, 3C, 3D, 4B, S3A and S3B; chi square for deviations from 50: 50 ratio in Figures 3E, 5A, 5B, 6C and S1C, and for deviations from 1 in Figures 1C and 4C.

## QUANTIFICATION AND STATISTICAL ANALYSIS

Statistical details of the experiments can be found in the figure legends and the Method Details section of the STAR Methods. Statistical details include statistical methods, exact value of n, what n represents and definition of center. Statistical tests were performed as described in the Method Details section using GraphPad Prism, and a *P* value of less than 0.05 was judged as statistically significant.

## Supplementary legends

**Figure S1. Biased flipping underlies the biased orientation of selfish centromeres towards the egg pole. Related to Figure 1.** (**A**) Schematic of female meiosis. The meiosis I (MI) spindle initially forms in the center of the oocyte and later migrates towards the cortex and orients perpendicular to the cortex, which is essential for the highly asymmetric cell division. (**B**) Schematic showing spindle asymmetry and biased orientation of larger centromeres in the intraspecific CHPO hybrid, based on previous results (Akera et al., 2017). Initial MT attachments are established when the spindle is still in the center and symmetric. Hybrid bivalents are off-center on the spindle, with the larger centromere closer to the pole, indicating that larger and smaller centromeres interact differentially with spindle MTs. Bivalent orientation on the spindle is unbiased right after spindle migration (Early Meta I), but the attachment of larger centromeres to the cortical side of the spindle is especially unstable, leading to detachment and flipping to establish biased orientation (Late Meta I). (**C**) CF-1 x CHPO (L x S) hybrid oocytes expressing CENP-B-mCherry and H2B-EGFP were imaged live after spindle migration. Time-lapse images show examples of flipping events to face larger centromeres towards the egg (top) or cortical (bottom, 0 – 30 min) side. Images are maximum intensity z-projections showing all chromosomes (left), or optical slices magnified to show flipping events (timelapse). Orange and white arrows indicate larger and smaller centromeres, respectively. Scale bar, 10 μm. Percentages indicate the frequency of each case (n = 14 flipping events from 29 cells). \**P* < 0.005, indicating significant deviation from 50%. Flipping events that faced larger centromeres towards the cortical side were subsequently reversed (bottom, 30 – 60 min), demonstrating the difficulty for larger centromeres to remain attached to the cortical side.

**Figure S2. BUB1, H2ApT121, and SGO2 are symmetric across bivalents in control oocytes. Related to Figure 2.** CF-1 x CF-1 (L x L) oocytes were fixed at metaphase I and stained for BUB1, H2ApT121, or SGO2. Images are maximum intensity z-projections showing all chromosomes (left), or optical slices magnified to show single bivalents (right). Quantification is shown in Figure 2.

**Figure S3. BUB1 targeting cancels asymmetry in both MCAK and CPC. Related to Figure 3.** (**A** and **B**) CF-1 x CHPO oocytes expressing Major Sat-BUB1 were fixed at metaphase I and stained for MCAK (A) or pINCENP (B). Images are maximum intensity z-projections showing all chromosomes (left) or optical slices magnified to show single bivalents (right). Scale bar, 10 μm. Graphs show centromere signal intensities or centromere signal ratios, calculated as the brighter divided by the dimmer signal for each bivalent. Each dot represents a single centromere (A and B, bottom, n > 53 for each condition) or a single bivalent (B, top, n > 51 for each condition). Red line, mean; \**P* < 0.001.

**Figure S4. CPC is spread across the chromosomes in the interspecific *spretus* hybrid. Related to Figure 4.** C57BL/6J x SPRET/EiJ (L x *sp*) oocytes were fixed at metaphase I and stained for pINCENP and CREST. Images are maximum intensity z-projections showing all chromosomes (left) or an optical slice magnified to show a single bivalent (right). Scale bar, 10 μm.

**Figure S5. Spindle asymmetry in tyrosinated MTs in the interspecific *spretus* hybrid. Related to Figure 6.** C57BL/6J x SPRET/EiJ (L x *sp*) oocytes were fixed at metaphase I and stained for tyrosinated α-tubulin and β-tubulin. Images are maximum intensity z-projections showing the whole oocyte (left) or a magnified view of the spindle (right). Dashed line, cortex; scale bars, 10 μm.

